# Integrative COVID-19 Biological Network Inference with Probabilistic Core Decomposition

**DOI:** 10.1101/2021.06.23.449535

**Authors:** Yang Guo, Fatemeh Esfahani, Xiaojian Shao, Venkatesh Srinivasan, Alex Thomo, Li Xing, Xuekui Zhang

## Abstract

The SARS-CoV-2 coronavirus is responsible for millions of deaths around the world. To help contribute to the understanding of crucial knowledge and to further generate new hypotheses relevant to SARS-CoV-2 and human protein interactions, we make use of the information abundant Biomine probabilistic database and extend the experimentally identified SARS-CoV-2-human protein-protein interaction (PPI) network *in silico*. We generate an extended network by integrating information from the Biomine database, the PPI network, and other experimentally validated results. To generate novel hypotheses, we focus on the high-connectivity sub-communities that overlap most with the integrated experimentally validated results in the extended network. Therefore, we propose a new data analysis pipeline that can efficiently compute core decomposition on the extended network and identify dense subgraphs. We then evaluate the identified dense subgraph and the generated hypotheses in three contexts: literature validation for uncovered virus targeting genes and proteins, gene function enrichment analysis on subgraphs, and literature support on drug repurposing for identified tissues and diseases related to COVID-19. The majority types of the generated hypotheses are proteins with their encoding genes and we rank them by sorting their connections to the integrated experimentally validated nodes. In addition, we compile a comprehensive list of novel genes, and proteins potentially related to COVID-19, as well as novel diseases which might be comorbidities. Together with the generated hypotheses, our results provide novel knowledge relevant to COVID-19 for further validation.

## Introduction

The COVID-19 pandemic, caused by severe acute respiratory syndrome coronavirus 2 (SARS-CoV-2), is a grave threat to public health and the global economy. It has led to more than 224 million confirmed cases and 4.62 million deaths worldwide as of September 12, 2021. SARS-CoV-2 is a newly discovered positive-sense single-stranded RNA virus and belongs to the member of the Coronaviridae (CoV) family [1]. It shares 89.1% and 50% nucleotide similarity to other previously detected human coronaviruses SARS-CoV and Middle East respiratory syndrome coronavirus (MERS-CoV) [2, 3, 4], respectively. Previous studies have also shown that they share stronger similarities with respect to their structures and pathogenicity [4]. These provide valuable knowledge to facilitate our understanding of the pathophysiology of SARS-CoV-2. Particularly, we now know that SARS-CoV-2 mainly enters human cells via binding of its spike protein to the angiotensin converting enzyme 2 (ACE2) receptor [5] and is associated with an extensive immune reaction referred to as “cytokine storm” triggered by the excessive production of interleukin 1 beta (IL-1b), interleukin 6 (IL-6), and others. However, much remains to be explored about how these critical human proteins are involved in the infection and the associated COVID-19 pathology [2], critical towards devising therapeutic strategies to counteract SARS-CoV-2 infection.

In order to investigate the complications and comorbidities of SARS-CoV-2 and to facilitate the search for effective treatment, many studies have been conducted to investigate the host dependencies of the SARS-CoV-2 virus from a systems level. For example, Blanco-Melo performed a comparative transcriptional analysis of COVID-19 patients responding to SARS-CoV-2 and other respiratory viruses, that revealed reduced innate antiviral defenses coupled with exuberant inflammatory cytokine production as the defining and driving features of COVID-19 [6]. Bojkova et al. conducted proteomic analysis to identify the host cell pathways that are modulated by SARS-CoV-2 and showed that inhibition of these pathways prevents viral replication in human cells [7]. Gordon et al. systematically mapped the interaction landscape between SARS-CoV-2 proteins and human proteins using affinity-purification mass spectrometry [8]. They identified 332 high-confidence protein interactions between SARS-CoV-2 viral proteins and human proteins related to various complexes and biological processes (about 40% of human proteins identified to interact with SARS-CoV-2 were associated with endomembrane system or membrane vesicle trafficking). From the presented SARS-CoV-2-human protein-protein interaction (PPI) network the authors identified 62 druggable SARS-CoV-2-interacting human proteins with 69 targeting ligands (drugs). Wei et al. [9] recently conducted genome-wide CRISPR screening and experimentally identified lists of pro/anti-SARS-CoV-2 genes. Stukalov et al. [10] profiled the interactomes of SARS-CoV and SARS-CoV-2 using A549 lung carcinoma cells. Li et al. [11] used genome-wide proteomic screening to identify cellular proteins that interact with SARS-CoV-2 proteins. The work by Gordon et al., Stukalov et al., and Li et al. have been reviewed in [12]. All these studies contributed to a better understanding of the SARS-CoV-2 and host protein interactome, providing insights for the development of therapies for the treatment of COVID-19. However, these studies only revealed different aspects of the potential mechanisms behind SARS-CoV-2 infection at specific conditions and do not bring out into open the comorbidities-, cell- and organ-type-specific human-viral interaction architecture.

To generate new knowledge, various computational approaches were applied to integrate the above experimental results with other information. Perrin-Cocon et al. [13] selected 112 publications including [8] that explicitly reported physical interactions between coronavirus viral and host proteins and assembled a coronavirus-host interactome. Krämer et al. [14] started from the PPI network identified by Gordon et al. and constructed 70 hypothesis networks using a machine learning algorithm. Gysi et al. [15] conducted research on identifying possible repurposing drugs for COVID-19 through predictive methods and they also retrieved PPIs from [8] and assembled a large human interactome during the process. Sadegh et al. [16] integrated SARS-CoV and SARS-CoV-2 virus-host interaction data from several sources including [8] and developed an online platform that implemented systems medicine algorithms for network-based prediction of drug candidates. As a common integration approach, integrative network analysis provides an efficient way to enable discovery and evaluation of (unknown) connections spanning multiple types of relationships inferred from different omics studies. Khorsand et al. [17] started from the most similar Alpha-influenzavirus proteins to SARS-CoV-2 proteins and constructed a weighted SARS-CoV-2-human PPI network. Messina et al. [18] constructed interactomes for SARS-CoV, MERS-CoV and HCoV-229E and employed random walk with restart (RWR) algorithm to identify the top 200 closest proteins for the selected three human coronaviruses. Zhou et al. [19] and Kumar et al. [20] have applied similar ideas to perform the integrative network analysis for elucidating the molecular mechanisms of SARS-CoV-2 pathogenesis.

To generate new hypotheses, we present a data analysis pipeline of integrative network analysis. Our work is different from most of the other methods, because we integrate experimentally validated results with a ‘large’ ‘probabilistic’ network. Analyzing ‘probabilistic’ networks increased the difficulty of the algorithm, but it can better incorporate the uncertainty in real-world. To make sense of a ‘large’ network, we need to focus on a small subset of important nodes and discard the rest of the nodes. In this work, we aim to identify high-connectivity sub-communities (connected to the experimentally validated results) from the integrated dataset since our fundamental assumption is that members in such sub-communities play more important roles in the network [20]. Specifically, in this work, we integrate the SARS-CoV-2 viral-human protein interaction (PPI) network [8], experimentally identified lists of pro/anti-SARS-CoV-2 genes [9], and a large probabilistic biological network called Biomine [21, 22]. Biomine integrates several databases including PubMed [23], UniProt [24], STRING [25], Entrez Gene [26, 23], and InterPro [27]. Many biological networks can be represented using probabilistic graph structures due to the intrinsic uncertainty present in their measurements. For instance, the edges in PPI networks obtained through laboratory experiments are often prone to measurement errors. The edges are often labeled with uncertainty levels that can be interpreted as probabilities. We aim to mine these probabilistic graphs to enhance our understanding of the SARS-CoV-2 and human protein interactions and to further aid the discovery of the essential/unknown knowledge relevant to the interactions between hosts and SARS-CoV-2 virus. The crux of our approach is to use a *core decomposition* strategy that detects highly connected sub-communities. Unlike other notions of cohesive subgraphs, e.g. *cliques*, *n-cliques*, *k-plexes*, which are all **NP**-hard to compute, *k-core* can be computed in polynomial time [28, 29, 30]. The goal of *k-core* computation is to identify the largest induced subgraph of a graph *G* in which each vertex is connected to at least *k* other vertices. The set of all *k-core*s of *G* forms the core decomposition of *G* [31]. The coreness (core number) of a vertex *v* in *G* is defined as the maximum *k* such that there is a *k-core* of *G* containing *v*. For an example of core decomposition, see Figure 1. For probabilistic graphs, the notion of core decomposition evolves into the more challenging probabilistic core decomposition. In this study, we connect the Biomine network to the small PPI network identified by Gordon et al. [8] and we integrate our previously proposed graph peeling algorithm [30] for probabilistic core decomposition and proposed an analysis pipeline to detect the probabilistic coreness in data, finding the high-connectivity sub-communities, and generate hypotheses on COVID-19 relevant bio-networks. The proposed analysis pipeline also supports integrating other experimentally validated results.

**Fig. 1.**
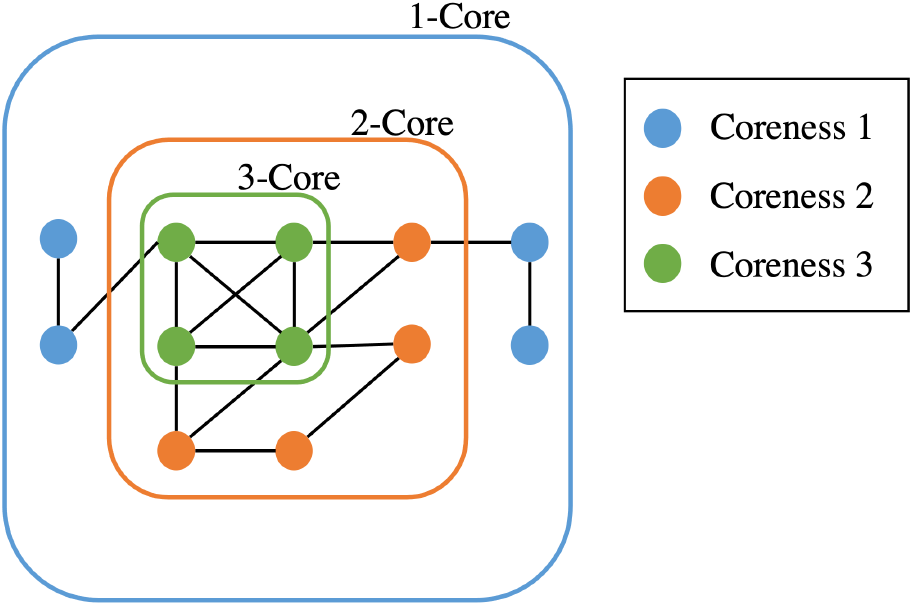
Core decomposition for an example graph

Specifically, from the results, we are particularly interested in dense cores (cores with high core number) in the Biomine database that overlap with the PPI network (and other integrated experimentally validated results) as much as possible. The dense cores in a different region of the Biomine network that do not overlap with the experimentally validated results are not of interest to us.

In short, our contributions are as follows.

1. **Pipeline on *k-core* decomposition and bio-network analysis**. A two-stage data screening procedure was added to the peeling algorithm (PA), making it focus more on cores with higher density. Overall, the pipeline consists of three steps: data preprocessing, PA, and functional enrichment analysis on the generated networks.
2. **Evaluation through literature mining and gene set enrichment analysis**. We evaluated the extended COVID-19 biological network in three contexts: literature support for identified tissues and diseases related to COVID-19, literature validation for uncovered SARS-CoV-2 targeting genes and proteins, and gene ontology (GO) over-representation test on the selected network for biological processes linked with RNA processing and viral transcription.
3. **COVID-19 hypotheses generation**. We discovered novel diseases that might be comorbidities, genes, and proteins that could potentially relate to COVID-19 and we presented them as top candidates for future validation.

## Methods and materials

### Biomine database and SARS-CoV-2-host protein-protein interaction network

The Biomine database is a large probabilistic biological network constructed using selected publicly available databases, for example, Entrez Gene, UniProt, STRING, InterPro, PubMed, Gene Ontology (GO), etc. The full Biomine database has 1508587 nodes, 32761889 edges and contains biology information of several species including humans. The SARS-CoV-2-host protein-protein interaction (PPI) network, identified by Gordon et al. [8], contains 360 nodes (27 SARS-CoV-2 viral proteins and 333 human proteins ^1^) and 695 edges. Since we are working with Homo sapiens data, as a preliminary stage of data screening, we select a subset of the full Biomine database, the *human* organism as the database to be used to extend the PPI network. This will eliminate approximately 43% of the full Biomine database. The *human* Biomine database contains 861812 nodes, 8666287 edges, and each entry possesses the form ***ε*** = (*from, to, relationship, link_goodness*). Here, *from* and *to* are two nodes forming an edge in the network, *relationship* is the link type describing the relationship between the two nodes, *link_goodness* is computed based on relevance, informativeness, and reliability [21] and is interpreted as the probability that the edge exists. A small sample from the *human* Biomine database is presented in Table S1 in Supplementary Methods and Results.

We then extend the PPI network with the *human* Biomine database and deal with duplicated entries and loops. For a detailed overview of the extension of SARS-CoV-2 and host protein-protein interaction network as well as duplicates/loops removal illustration, see relevant section and Figure S1 in Supplementary Methods and Results. As mentioned before, since Wei et al. [9], Stukalov et al. [10] and Li et al. [11] also recently experimentally validated lists of pro/anti-SARS-CoV-2 genes and viral-host PPI networks, we integrate the findings of [9] in our research and we retain [10, 11] for validations. Details of this analysis can be found in later sections including the data analysis pipeline section and validation from independent evidence section.

### Definition of probabilistic core decomposition

Consider a probabilistic graph 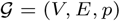 where *V* is the set of vertices, *E* is the set of edges, and *p* is a mapping function that aligns each edge *e* ∈ *E* to its existence probability *p*_*e*_. Given a vertex *v* ∈ *V*, let *d*_*v*_ be the number of edges incident on *v*. Each *possible world* 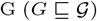, is a deterministic version of 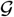 in which a subset *E*_*G*_ ⊆ *E* of edges appear. Let 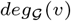 be a random variable with values in [0, *d*_*v*_], and distribution:

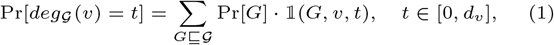

where 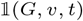 is an indicator function which takes on 1 if *v* has degree equal to *t* in the possible world *G*. *η-degree* of vertex *v*, denoted by *η-deg*(*v*), where *η* ∈ [0, 1], is defined as the highest *t* ∈ [0, *d*_*v*_] for which 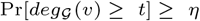 [30, 28].

For the (*k, η*)-core of probabilistic graph 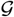, we use the definition given by [28]: the (*k, η*)-core of probabilistic graph 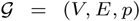 is the maximal induced subgraph 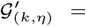 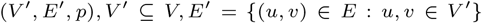 where the *η-deg*(*v*) for each *v* ∈ *V′* is at least *k*. The largest *k* for which *v* is a part of a (*k, η*)-core is called *η*-core number or probabilistic coreness of *v*. The set of all (*k, η*)-cores is the unique core decomposition of 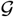 and follows the following relation [30]:

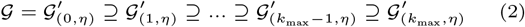

where *k*_max_ is the maximum probabilistic coreness of any vertex in 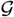. For simplicity, we will use core number and coreness instead of *η*-core number/probabilistic coreness in the rest of the paper. Additionally, we use *dense* to describe cores with high core numbers.

### Data analysis pipeline

The Biomine database contains abundant biological information. Though we used the smaller *human* Biomine database, after extending it with the SARS-CoV-2-host protein-protein interaction (PPI) network the resulting network is still enormous and hard to reason. To make sense of such a huge network, we propose a data analysis pipeline with three steps described below. An illustration of the workflow of data processing and analysis pipeline can also be found in Figure 2. Pseudocode with more details of the data analysis pipeline can be found in Algorithm S1 in Supplementary Methods and Results.

**Fig. 2.**
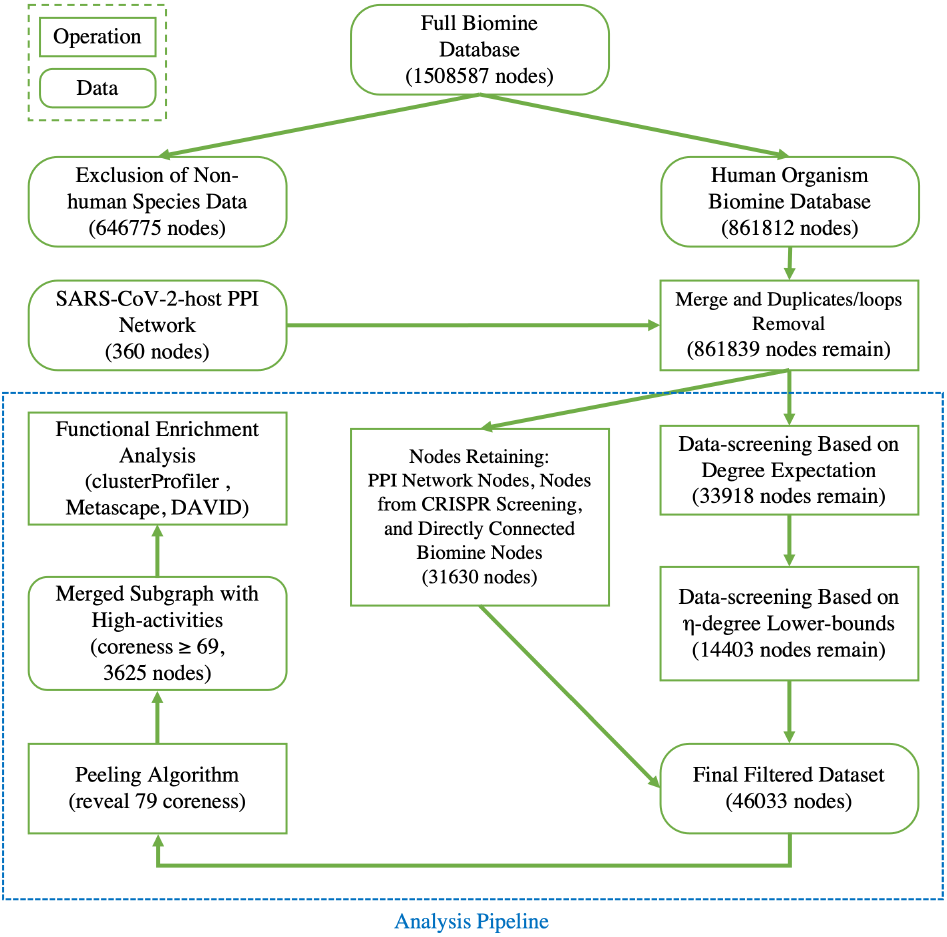
Workflow of data processing and analysis pipeline

### Step 1: Data preprocessing

The data preprocessing step contains three sub-steps, screening based on degree expectation, screening based on lower-bounds of *η-degree*, and nodes retaining. The goal of this data preprocessing step is to reduce the nodes in the network and speed up the follow-up analyses. Specifically, we remove large proportions of nodes with too low connectivity (i.e. vertices with small *η-degree*) to be members of dense sub-communities in the network.

Here we start with the first sub-step and we briefly explain the methodology and the rationale behind it. For each vertex *v* ∈ *V*, we have a set of edges incident to *v* and each edge is accompanied with a probability of existence *p*_*i*_ that is independent of other edge probabilities in 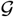. Given a probabilistic graph 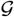, and a vertex *v*, 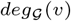 can be interpreted as the sum of a set of independent Bernoulli random variables *X*_*i*_’s with different success probabilities *p*_*i*_’s [30] where:

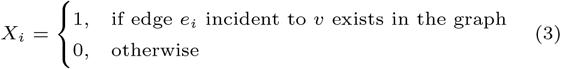

and 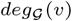 follows Poisson binomial distribution with 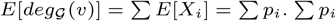. ∑*p*_*i*_ therefore can be seen as an approximation to 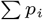 so we use ∑*p*_*i*_ as the first screening criteria. As thresholds are user-defined, any positive integer greater or equal to 0 is accepted but it is recommended that the first threshold is larger than 5 (if the first threshold is set to 5 it means any nodes with an expectation of degree less than 5 is removed, e.g. only nodes with ∑*p*_*i*_ ≥ 5 are kept). The goal of this step is to remove nodes that are rarely connected with others and hence are not eligible to be part of any highly connected sub-network. For example, if a vertex *u* has ∑*p*_*i*_ less than 5, it means that 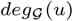 will also likely be around 5 with slight variations, and hence *u* will not appear in cores with high activities (e.g. vertices with high coreness). If the first threshold is set lower, more nodes are retained. To speed up subsequent analyses, the threshold value should be high enough, yet it cannot be so high that possible highly connected nodes are removed. In our experiment, we empirically choose a conservative number, 5, as the first threshold, but other threshold values could be used.

For our merged dataset, 33918 nodes passed the screening. Now, we introduce the second step of data prepossessing.

For the nodes that passed the first stage of data screening, we calculate lower-bounds of their *η-degree* using Lyapunov Central Limit Theorem (CLT) implemented in PA. We select 10 as the passing threshold (e.g. if a node has *η-degree* lower-bound greater than or equal to 10, we keep it, otherwise it will be removed). The rationale is if a vertex has at least 10 edges incident to it before peeling, we can consider it a hotspot suited for the afterward high activity subgraph mining. If in the full network a node is not connected to at least 10 other nodes, there is no point in performing core decomposition as we only focus on dense sub-communities. Note that as we start the peeling process, the node’s *η-degree* will also start decreasing, so in this last data filtering stage, we only select nodes based on their initial *η-degree* lower-bound.

There are many nodes in Biomine that can be directly connected to the PPI network, which are potentially more useful than other nodes. In the nodes retaining step, we force them to not be screened out and let downstream analyses decide whether they are useful.

Besides the PPI network, we also treat the list of experimentally validated genes [9] as ground truth and retain their mapped nodes for downstream analysis without the data screening steps. Wei et al. [9] selected the top and bottom 250 genes from their sorted gene list for analysis so we also select these 500 genes in our research. We map the selected list of genes to its corresponding UniProt indexes in Biomine. When UniPort index is not available, STRING index is used and those that still failed to map are discarded. Eventually, 439 of the genes are mapped and integrated into our research. There is an overlap of 8 nodes between the PPI network by Gordon et al. and the CRISPR screened gene list by Wei et al. after mapping. In total, we obtained 791 experimentally validated ground truth nodes.

In the merged dataset, 30839 unique Biomine nodes are directly connected to the nodes in the PPI network and nodes in the selected CRISPR screened gene list (a total of 31630 nodes). To preserve valuable information, we retain them from data screening. All other nodes in the *human* Biomine database that are not directly connected to the ground truth nodes will be subject to the two data screening steps.

After this step, a total of 46033 nodes remained in our dataset and the filtered dataset was passed to PA for probabilistic core decomposition with *η* set to 0.5. For a discussion on different settings of *η*, see Discussion section.

### Step 2: Peeling algorithm to find coreness of nodes

In this section, we briefly describe the graph peeling algorithm (PA) introduced by [30].

As mentioned before, *η-deg*(*v*) is defined as the highest *t* for which 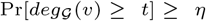. In [30], Lyapunov Central Limit Theorem (CLT) is applied to approximate 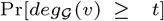 and find the largest *t* such that 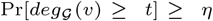. They showed that Lyapunov CLT can produce a very accurate lower-bound on vertex’s true *η-degree*. Since the lower-bound is easy to compute, it helps reduce PA’s running time significantly.

To summarize PA, it first computes the lower-bound on the *η-degree* for each vertex using Lyapunov CLT, then it stores vertices in an array in ascending order of their lower-bound values. Then, the algorithm starts processing vertices based on their (lower-bound on) *η-degree*. When a vertex *v* is being processed, PA algorithm determines whether *v*’s exact *η-degree* is available or *v* is on its lower-bound. If the former criteria holds, PA sets *v*’s coreness to be equal to its *η-degree* at the time of process, removes *v*, and decreases the *η-degree* (exact or lower-bound) of *v*’s neighbours by one. Otherwise, if *v* is on its lower-bound, *v*’s exact *η-degree* is computed and *v* is swapped to the proper place in the array. At the end, (*k, η*)-core of 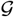 is obtained by collecting all the vertices with coreness at least *k*. Note that for efficiency reasons, (1) lower-bounds are used in the main parts of PA where *η-degree* values are required, and only when a vertex becomes a candidate for removal the exact *η-degree* is calculated, (2) after removing a vertex *v*, the step of updating the *η-degree* of *v*’s neighbours is delayed as much as possible (*lazy updates* strategy).

### Step 3: Functional enrichment analysis

Pathway analysis could prove crucial in understanding how the virus infects the human body [7]. To evaluate functional pathways of proteins involved in SARS-CoV-2 host interactions from the core decomposition result of PA, gene enrichment analysis was performed using clusterProfiler [32] and Metascape [33].

P-values were calculated by hypergeometric test [34], adjusted using Benjamini–Hochberg procedure [35], and adjusted p-value *<* 0.01 were used as the threshold of significance to control the false-discovery rate. We also performed DAVID functional annotation clustering [36, 37] on selected subgraphs. Since Metascape and DAVID both restrict input gene list size up to 3000, if our list exceeds that number, we will select the top 3000 nodes based on their connections to the integrated experimentally validated nodes. For UniProt indexing nodes, we select UNIPROT_ACCESSION as the gene list identifier. Everything else is kept as default.

## Results

We merged the original SARS-CoV-2-host protein-protein interaction (PPI) network [8] with the *human* Biomine database and we removed duplicated and looped edges in the merged dataset. We also integrated the experimentally validated pro/anti-SARS-CoV-2 gene lists by Wei et al. [9]. We then passed the merged dataset through our proposed analysis pipeline. Approximately 5.3% of nodes remained after data screening and the algorithm revealed the presence of 88 cores. More specifically, the nodes that remained in the filtered dataset yielded 79 different coreness values ranging from 0 to 88. Many node types exist in the filtered dataset, for example, UniProt, STRING, PubMed, GO (including indexes for biological process, cellular component, molecular function), etc. Approximately 11.35% of nodes were assigned coreness 1 and 2, which accounts for 2685 nodes and 2539 nodes, respectively. The 791 experimentally validated nodes (including the original SARS-CoV-2-human protein-protein interaction (PPI) network and the pro/anti-SARS-CoV-2 gene lists by Wei et al.) were distributed across 74 different coreness with minimum coreness of 1 and maximum coreness of 88. The nodes that were directly connected to the experimentally validated nodes (we refer to them as *level 1 connections*) had similar node count distribution with roughly 16.88% nodes assigned with coreness 1 and 2, followed by coreness 77 that contains 2108 (≈ 6.84%) nodes.

Table 1 shows the top-10 coreness values in terms of node count for the three scenarios: experimentally validated nodes, level 1 connections, and complete nodes set. As can be seen from Table 1, core 69 and core 77 are the two most frequently appeared denser cores that contain a significant fraction of nodes. As indicated before in Equation 2, the denser cores are at the same time among the smaller cores, so we further merged all cores that are denser than 68 (i.e. coreness 69, 70, 71, 72, 73, 74, 75, 76, 77, 88) into a giant subgraph to avoid losing potential connections as well as the corresponding information between cores.

**Table 1.**
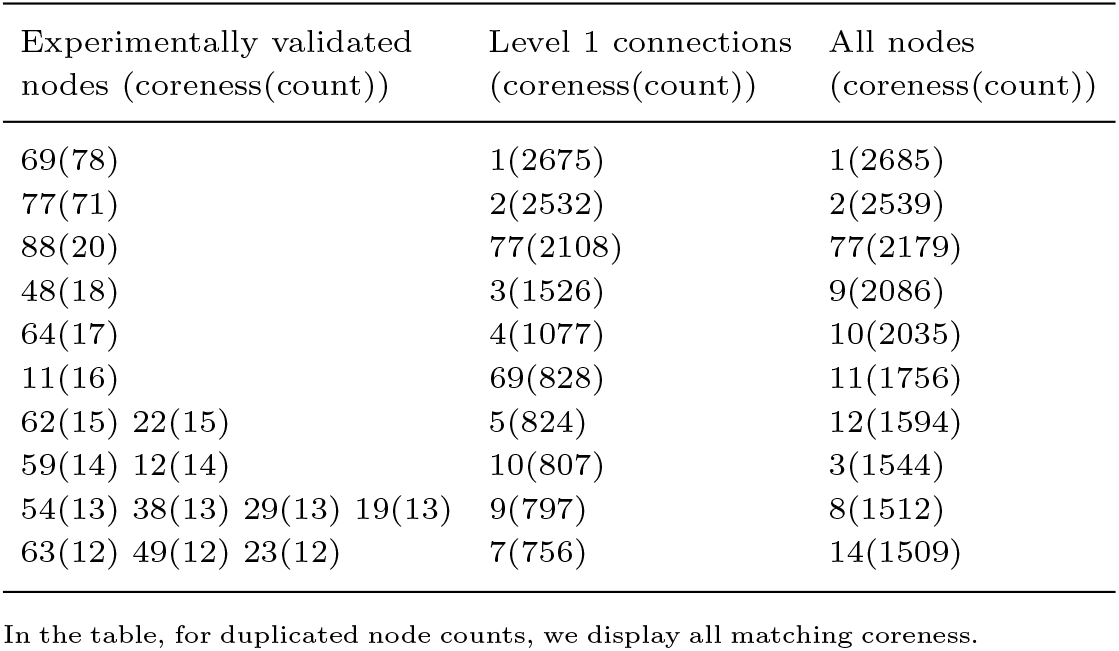
Top-10 coreness values revealed by peeling algorithm (sorted by node count)

The resulting subgraph contains 3625 nodes and 2025378 edges. By definition, the merged subgraph is the same as core 69 (it contains nodes with coreness 69 and above). Among these 3625 nodes, a total of 179 nodes are experimentally validated and 57 (≈ 17.12%) SARS-CoV-2 interacting human proteins identified by Gordon et al. [8] can be found in the subgraph. The other 3446 nodes are all level 1 connections. In addition, a majority of nodes (3485, ≈ 96.14%) in the subgraph are proteins labeled with corresponding UniProt ID. A complete dissection of node types for the subgraph can be found in Table S2 in Supplementary Methods and Results.

### SARS-CoV-2 associating genes discovery

We hypothesized that other protein nodes connected with the reported 57 SARS-CoV-2 interacting human proteins in different cores within the subgraph may contain potential missing connections from the single experiment, and provide novel molecular components for better understanding the pathogenicity of SARS-CoV-2 infection which will eventually be beneficial for identifying new biological/pharmaceutical targets.

We first explored the distribution of the 57 SARS-CoV-2 interacting human proteins in the merged subgraph, and observed that 27 of them has coreness 69, 2 of them has coreness 71, 1 of them has coreness 73, 20 of them has coreness 77, and 7 belongs to core 88 with coreness 88 assigned (Table 2). Since our goal is to extend the integrated experimentally validated results (we mainly want to focus on the SARS-CoV-2-human protein-protein interaction (PPI) network found by Gordon et al.) and generate more research hypotheses, in the merged subgraph (the extended network), all the nodes not belonging to the original PPI network or the pro/anti-SARS-CoV-2 gene lists can be considered as generated hypotheses. We then try to identify the hypotheses that had already been studied by other researchers. From the subgraph, we obtained 3306 UniProt indexing nodes that are directly connected to the 179 experimentally validated nodes. We further ranked these 3306 nodes by their connections to the reported experimentally validated results (we will refer to this number as *nConnect*, the complete ranked list can be found in Supplementary Table 5) and performed literature mining on their associations/correlations with COVID-19. After an automatic web crawling followed by manual inspection, we identified 306 nodes (≈ 9.26%) that show associations or correlations with COVID-19 supported by at least one study. Among the 306 literature verified nodes, 68 of them has coreness 69, 193 of them has coreness 77 (in total, ≈ 85. 29% of literature verified nodes were either assigned coreness 69 or coreness 77, a complete plot on coreness distribution is shown in Figure S2 in Supplementary Methods and Results). As mentioned before, core 69 and 77 are most frequently appeared and showed denser sub-communities compares with others in different contexts in Table 1. And for the 57 SARS-CoV-2 interacting human proteins in core 69, 47 of them (≈ 82.46%) either have coreness 69 or coreness 77. We believe these findings support our assumption that more valuable information can be found in dense sub-communities and we move on to explore the criteria for high quality hypothesis (i.e. hypothesis that most likely to be true).

**Table 2.** Node distribution of the merged subgraph

| Coreness | Total number of nodes | Total number of PPI network nodes |
| --- | --- | --- |
| 69 | 906 | 27 |
| 70 | 56 | 0 |
| 71 | 61 | 2 |
| 72 | 46 | 0 |
| 73 | 54 | 1 |
| 74 | 10 | 0 |
| 75 | 21 | 0 |
| 76 | 38 | 0 |
| 77 | 2179 | 20 |
| 88 | 254 | 7 |

Table 3 lists the top genes and encoded proteins that are connected to more than half of the integrated experimentally validated nodes in the subgraph, ranked by *nConnect*. Over 20% of nodes in Table 3 received literature supports. Therefore, we consider nodes in Table 3 to be high quality hypotheses compared to the rest of the nodes in the subgraph and we recommend researchers start validating them first. The subgraph node list and the complete list of nodes that received literature supports can be found in Supplementary Tables 1 and 2. A detailed discussion on the literature-supported nodes in Table 3 can be found in the Discussion section.

**Table 3.**
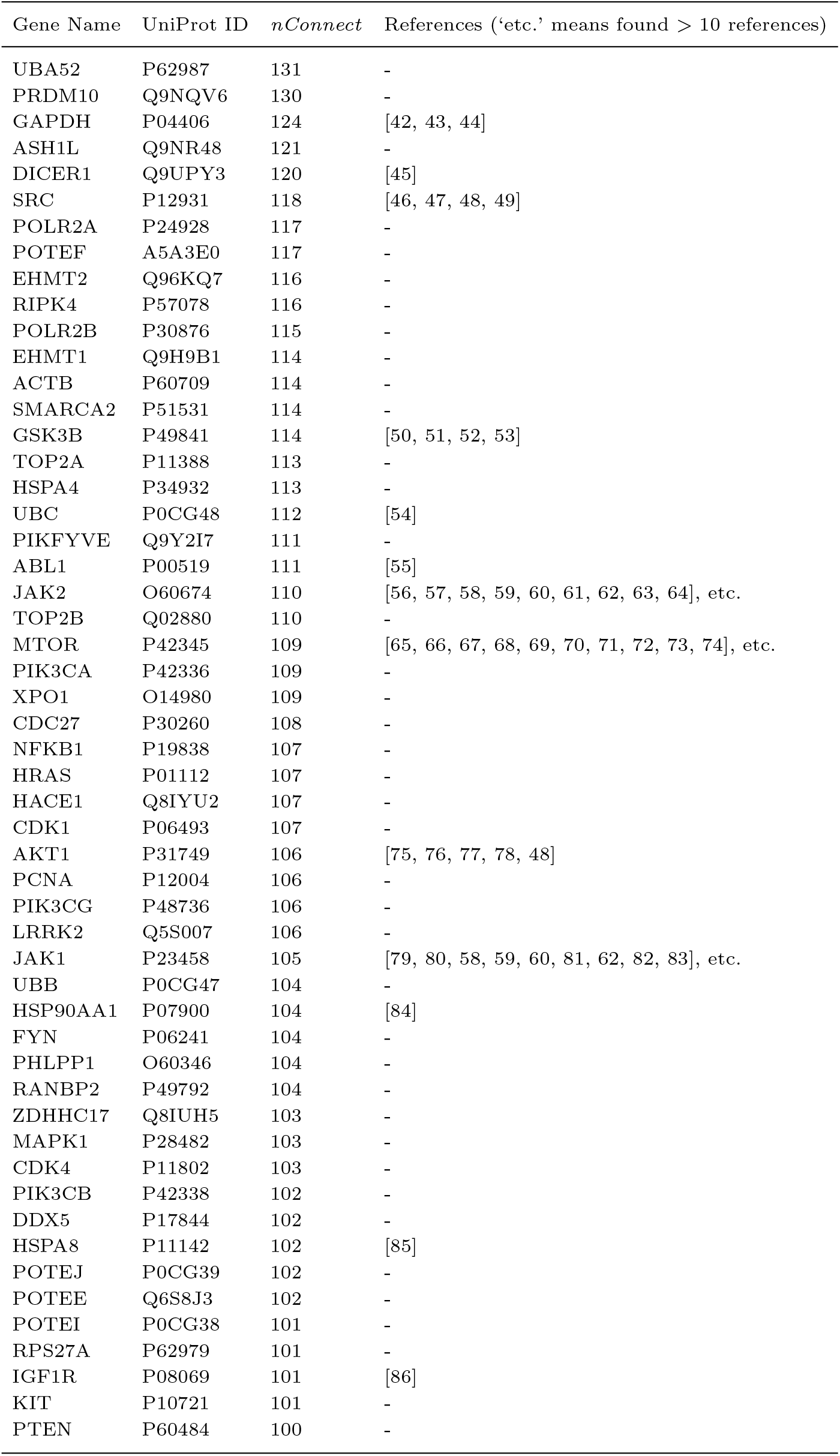
Discovered top genes potentially related to COVID-19

### Gene ontology over-representation analysis

We first performed an enrichment test of gene ontology (GO) biological process for genes that encode all of the UniProt indexing nodes in the subgraph using the enrichGO function of clusterProfiler package in R with default parameters [32]. The top 30 GO terms with the smallest adjusted p-value were presented in Figure 3. The most significant GO term is *protein polyubiquitination* (adjusted p-value ≈ 8. 96 * 10^−75^), which accounts for 204 of the total 3352 mapped input genes (≈ 6.09%), followed by *ribonucleoprotein complex biogenesis* (adjusted p-value ≈ 2. 42 * 10^−58^). The top enriched term might suggest that SARS-CoV-2 hijacks cell’s ubiquitination pathways for replication and pathogenesis, which is one of the findings of [8]. Interestingly, the virus-associated biological processes *viral gene expression* and *viral transcription* were also found to be enriched and account for about 3.73% and 3.49% of input eligible gene set, respectively. In addition, we noted that 10 out of the top 30 GO terms were RNA-related. For example, *mRNA catabolic process* (6.06% of total input eligible genes), *RNA catabolic process* (6.38%), and *nuclear-transcribed mRNA catabolic process, nonsense-mediated decay* (3.04%) were the top items. This is in line with Gordon’s study where they found SARS-CoV-2 proteins NSP8 and N involved in RNA processing and regulation [8].

**Fig. 3.**
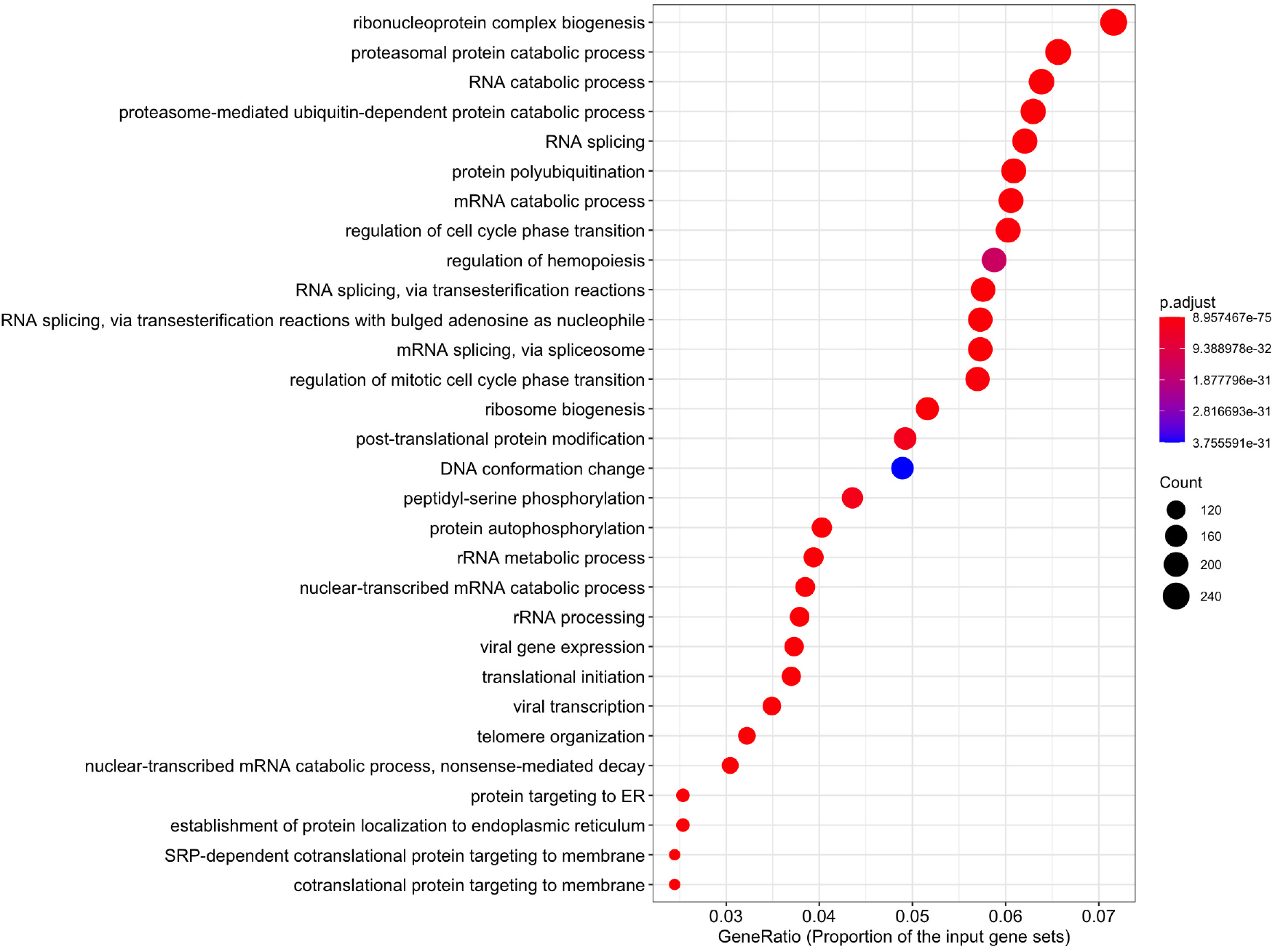
Top GO terms of all mapped UniProt indexing nodes in the subgraph ranked by GeneRatio

We further used Metascape v3.5-210201 [33] to perform pathway and process enrichment analysis on UniProt indexing nodes with different coreness. Two subsets showed significant enrichment in the molecular functions of immune response to bacteria or viruses. Specifically, 5.77% protein-encoding genes with coreness 70 enriched the term *The human immune response to tuberculosis* and other 5.77% genes enriched the term *regulation of viral process*, see Figure 4-(a) for other relevant enriched items (*negative regulation of protein kinase activity*, *Signaling by Receptor Tyrosine Kinases*, etc.). Regarding UniProt indexing nodes with coreness 73, network enrichment analysis using Metascape returned a few significant modules (Figure 4 -(b)). Interestingly, G2/M DNA damage checkpoint is found to be the top enriched term which is consistent with the observations from recent studies. For instance, Garcia Jr. et al. [38] detected key proteins involved in cellular signaling pathways mTOR-PI3K-AKT, ABL-BCR/MAPK and DNA-damage response that are critical for SARS-CoV-2 infection, and further they identified DNA-damage response inhibitor as potent blocker of SARS-CoV-2 replication. Bouhaddou et al. [39] infected Vero E6 cells with SARS-CoV-2 and observed a significant increase in the fraction of cells at the G2/M transition phase. Gordon et al. [8] found that SARS-CoV-2 NSP1 protein interacts with all four members of the DNA polymerase alpha complex, which couples DNA replication with DNA-damage response [40]. Additionally, Blanco-Melo et al. [6] found the excessive expression of cytokine as one of the strong features of SARS-CoV-2 infection, here we are able to find several terms that are related to cytokine storm induced by SARS-CoV-2 (see Figure 4-(b) for details). For instance, *IL-4 Signaling Pathway* and *Cytokine Signaling in Immune system* are related to cytokine storm upon virus infection. Particularly, IL-4 is one kind of cytokine that acts as a regulator of the JAK-STAT pathway and contributes to human body immune responses [41].

**Fig. 4.**
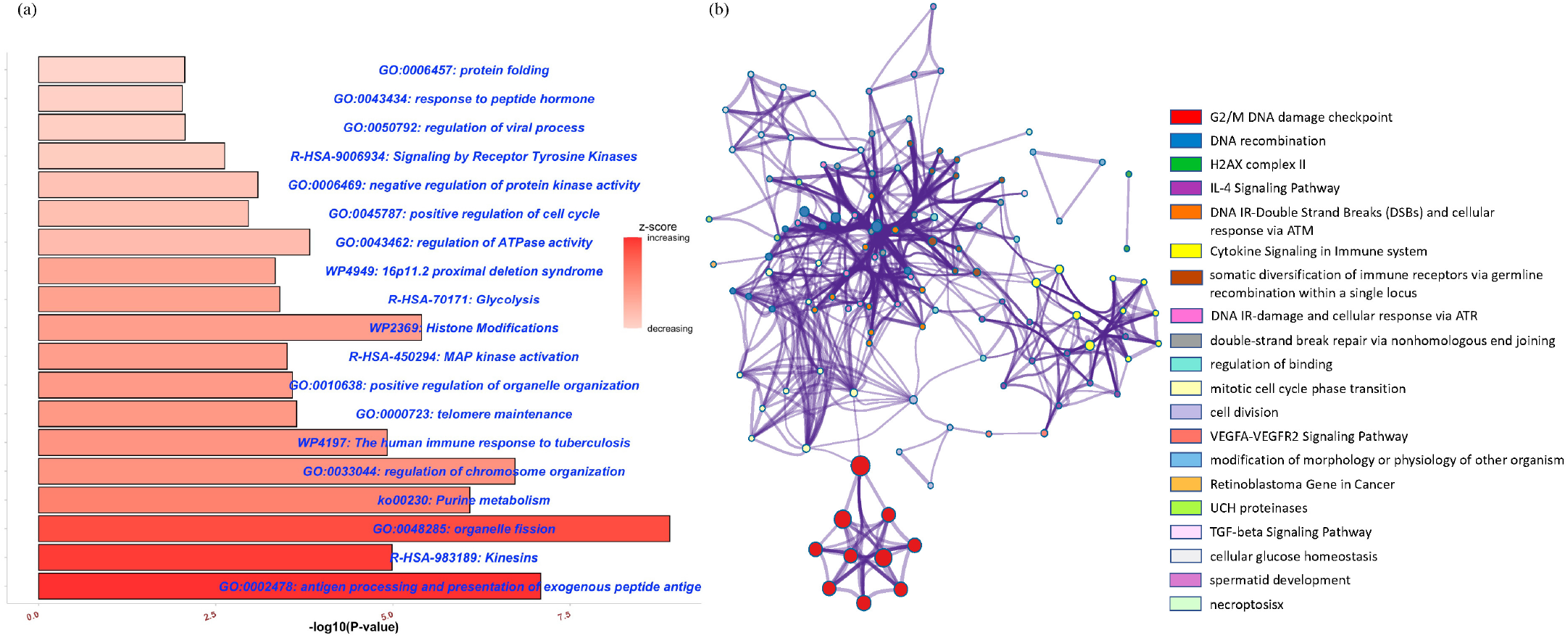
Metascape enrichment results for coreness 70 and 73. **(a)** Top biological terms (pathway, process, etc.) enriched for coreness 70 ranked by p-values **(b)** Network of enriched terms for coreness 73 colored by cluster ID, nodes shared the same cluster ID typically lie close together. Node size is proportional to p-value significance.

### SARS-CoV-2 interacts with tyrosine-related proteins

Bouhaddou et al. [39] found changes in activities for 97 out of 518 human kinases during SARS-CoV-2 virus infection.

Surprisingly, among the 97 kinases list they discovered, 76 were found in our merged subgraph (≈78.35%). This motivated us to check for all the 321 kinases-related UniProt indexing nodes in our subgraph (the complete list of kinases-related nodes can be found in Supplementary Table 3). Interestingly, 85 of them are tyrosine-related nodes (≈26.48%). As mentioned previously, we found 306 out of the 3306 UniProt indexing level 1 connections (≈ 9.26%) in the merged subgraph that had at least one study showing they have some relations with COVID-19. When we restrict to tyrosine-related proteins, this proportion increased more than threefold to 28.24% (Table 4). That is, 24 of the 85 tyrosine-related proteins (Supplementary Table 4) in the merged subgraph have been explored for their associations with SARS-CoV-2 virus, including SRC, JAK1, JAK2, IGF1R, ABL1, TYRO3, etc. The high proportion of validated tyrosine-related proteins further motivated us to perform a DAVID functional annotation clustering [36, 37] on level 1 connections UniProt indexing nodes in core 69 (the merged subgraph). In total, 341 clusters were identified and the term *Tyrosine-protein kinase* (fold-change = 5.1, p-value = 1. 5 * 10^−44^) is among the top 10 clusters ranked by enrichment score (the DAVID analysis report can be found in Supplementary Report 1). Of particular importance, SRC, JAK1, JAK2, ABL1 are among the top of the list with large connections to the integrated experimentally validated nodes (Supplementary Table 5). Since our main focus is the PPI network found by Gordon et al., in Figure 5 we presented a complete PPI network between tyrosine-protein kinase SRC, JAK1/2, ABL1/2 and their connections to the SARS-CoV-2 interacting human proteins in the subgraph. We also include the SARS-CoV-2 viral proteins (red diamond) from the original PPI network. This network covered half (n=13) of the 27 SARS-CoV-2 viral proteins, including envelope (E), NSP7, NSP9, ORF10, etc. It showed these five proteins were not directly interacting with the virus but at level one connections where SRC, JAK1/2, and ABL1/2 are hub nodes and connect with each other (connection degree being 42, 37, 37, 36, and 29, respectively).

**Table 4.**
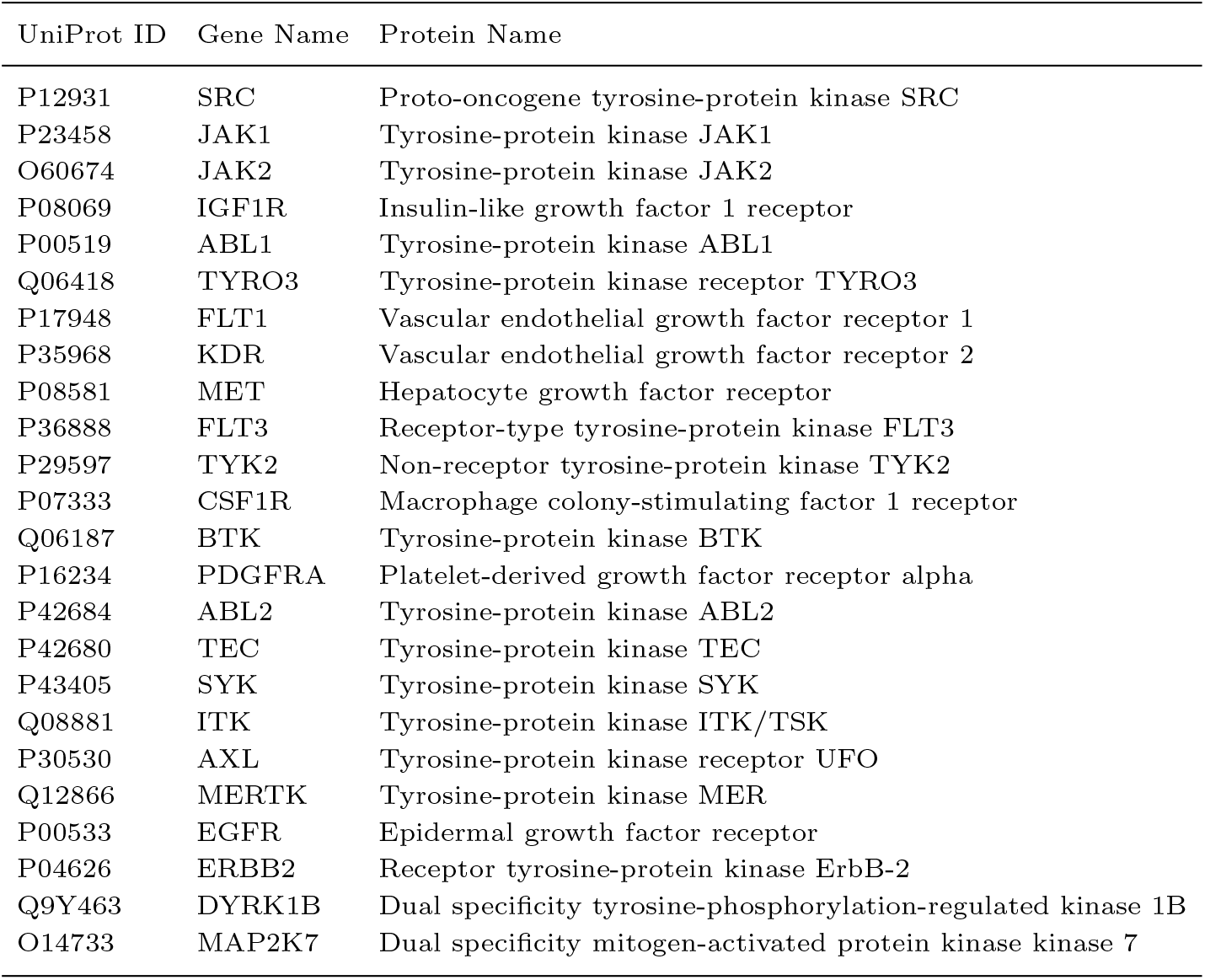
All tyrosine-related proteins in the subgraph that received literature support

**Fig. 5.**
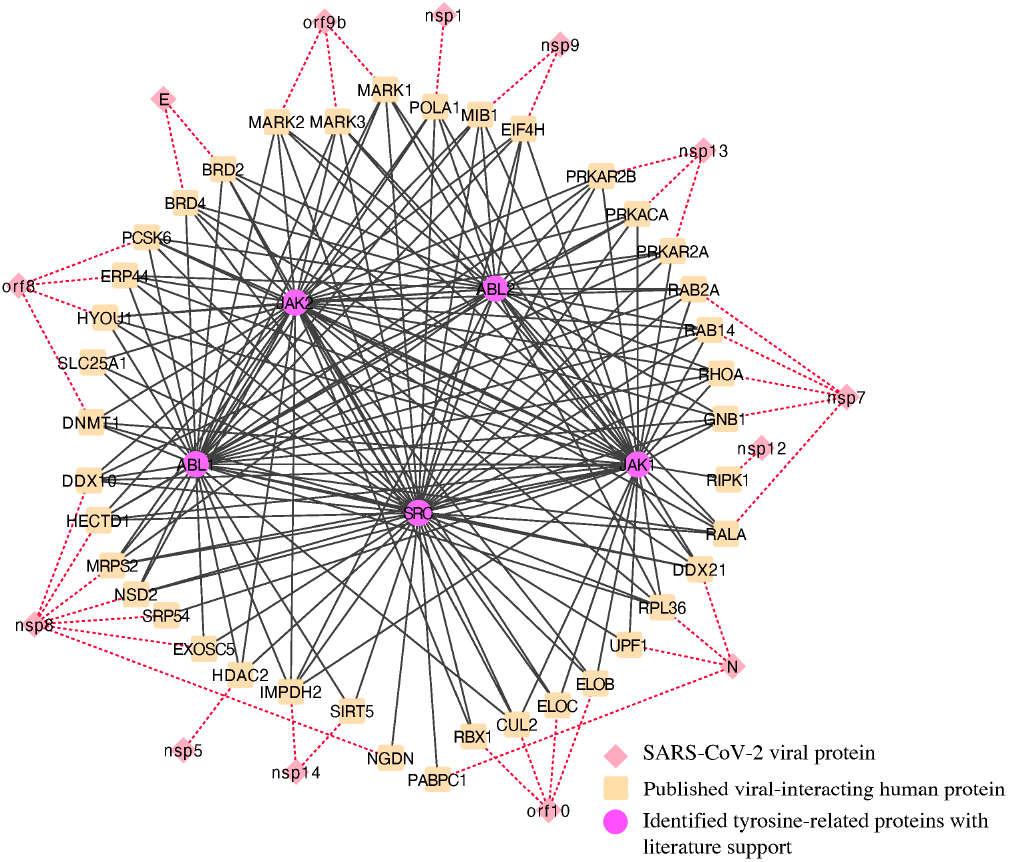
SARS-CoV-2 protein interaction with Tyrosine-protein kinase JAK1/2, ABL1/2, SRC. Dashed edges indicate the proteins do not belong to the identified subgraph

Additionally, we ranked all the nodes in the merged subgraph by their degrees and investigated the top 5% of the sorted nodes (181 out of 3625 nodes, Supplementary Table 1). There are 102 “kinase” protein nodes and 53 of them are “Tyrosine-protein kinase”. Among the 53 tyrosine-related nodes, the top ranking ones were SRC (ranked 5 out of 181), ABL1 (ranked tie at 19 out of 181), and JAK2 (ranked 22 out of 181) with degree connection 2687, 2483, and 2477, respectively. Take together, evidence obtained so far highly implied that SRC, ABL1, and JAK2 are three important hub genes.

### COVID-19 related tissues and diseases discovery

Through the 332 human viral-interacting proteins across different tissues, Gordon et al. [8] identified lung as the tissue with the highest level of differential expression of the SARS-CoV-2 interacting human proteins. They also found 15 other tissues with a high abundance of SARS-CoV-2 human interacting proteins. In our subgraph, we were able to locate 5 out of the top 16 tissues identified by [8]: lung (coreness=69), testis (coreness=69), placenta (coreness=69), liver (coreness=69), and brain (coreness=77). Curiously, we also discovered two diseases that seem to be associated with COVID-19 within the subgraph (Supplementary Table 5): cervix carcinoma (coreness=70) and erythroleukemia (coreness=69). Figure 6-(a) presents a network consisting of the 5 identified tissue nodes, 2 disease nodes, and 2988 UniProt indexing nodes that directly connect to tissue nodes and disease nodes in the subgraph (an interactive version of the network can be found in our GitHub repository, see Supplementary materials). 54 out of the 57 reported SARS-CoV-2 interacting human proteins can be found in this network. Figure 6-(b) shows the interaction between the 54 SARS-CoV-2 interacting human proteins and the tissue and disease nodes identified in the subgraph under a higher resolution. Similar to what Gordon et al. [8] found, lung, testis, placenta, liver, and brain are heavily involved during SARS-CoV-2 infection. For example, the HDAC2 protein, which is observed to be expressed in testis, lung, overexpressed in cervical cancer, etc., has been identified by Gordon et al. to be associated with the SARS-CoV-2 NSP5 protein. We noted that the same SARS-CoV-2 interacting human proteins are also expressed in cervix carcinoma and erythroleukemia in the sub-graph. We hypothesized drugs used to treat these two diseases might be useful to treat COVID-19.

**Fig. 6.**
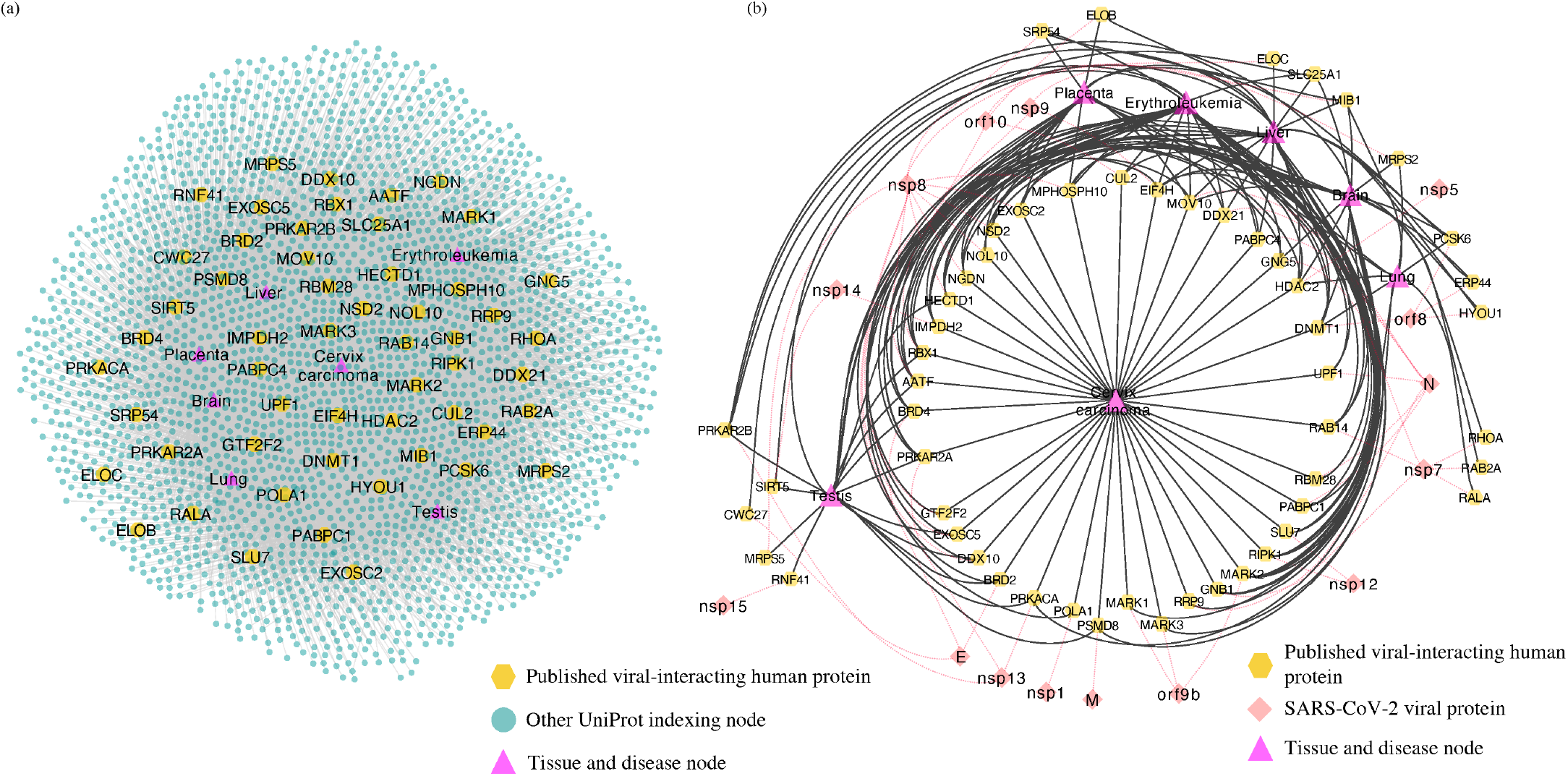
Interactions between tissue, disease, and UniProt indexing nodes in the subgraph. All of the edges have *is_expressed_in* as edge relationship type. **(a)** Interaction map between UniProt indexing nodes, identified tissue nodes and disease nodes **(b)** Association between SARS-CoV-2 host interacting proteins and tissue/disease nodes in core 69, we add SARS-CoV-2 viral proteins (red diamond with dashed links, indicating they do not exist in the subgraph) for clarity

Idarubicin, daunorubicin, and cytarabine are the three drugs used to treat erythroleukemia. Chandra et al. [87] suggest idarubicin as a potential drug that can be repurposed for controlling SARS-CoV-2 infection due to its good binding affinity to SARS-CoV-2 NSP15 encode endoribonuclease enzyme.

Bevacizumab is used to treat cervix carcinoma. According to Rosa et al. [88], there have been clinical trials about repositioning bevacizumab for COVID-19. In addition, Amawi et al. and Zhang et al. [89, 90] identified bevacizumab as one of the promising therapeutic treatments against COVID-19.

### Validation from independent evidence

To validate our generated hypotheses with independent evidence, we compared them with results obtained through both computational and experimental approaches. The consistent parts are considered as independent evidence of each other, whereas the non-overlap parts in our result can be considered as novel findings. It is worth noting that our work is the only one that starts with probabilistic networks and performed integrative network analysis.

First, to investigate how our results relate or overlap with the results obtained by other computational approaches, we compared our generated hypotheses with their presented SARS-CoV-2-related interactomes, proteins, and protein-protein interactions. Perrin-Cocon et al. [13] assembled a coronavirus-host interactome and 325 (28.63%) of the total 1135 human proteins mapped to UniProt database within the constructed network can be found in our merged subgraph. Krämer et al. [14] constructed 70 hypothesis networks using a machine learning algorithm and 197 (37.31%) of the 528 proteins mapped to Homo sapiens UniProt database can be found in our merged subgraph. Gysi et al. [15] searched extensively in the literature and public databases to assemble a large human interactome with the goal of identifying possible COVID-19 repurposing drugs. Their constructed interactome contains 17349 proteins that can be mapped to Homo sapiens UniProt database which results in a huge overlap (3320 proteins) with our merged subgraph. However, there are still 165 (4.74%) UniProt indexing nodes in our subgraph that they are not able to identify despite their extensive search in literature. Sadegh et al. [16] developed an online platform that implemented systems medicine algorithms for network-based prediction of drug candidates. Using the 332 viral-interacting human proteins identified by Gordon et al. [8] as input to their default Multi-Steiner algorithm for drug target discovery with default parameters we obtained a total of 393 human proteins, with 332 being the original human proteins in the input. In the rest of 61 proteins, 29 (47.54%) can be found in our merged subgraph. In addition, Khorsand et al. [17] constructed a three-layer SARS-CoV-2-human PPI network using Alpha-influenzavirus proteins that are most similar to SARS-CoV-2 proteins and 662 (34.88%) of the identified 1898 UniProt mapped human proteins in their network can be found in our merged subgraph. Messina et al. [18] constructed interactomes for three human coronaviruses: SARS-CoV, MERS-CoV and HCoV-229E and employed random walk with restart (RWR) algorithm to identify the top 200 closest proteins for each of the virus (total 600 proteins). 548 proteins can be mapped to Homo sapiens UniProt database and 241 (43.98%) of them can be found in our merged subgraph. In total, 3323 (95.35%) of the UniProt indexing nodes in our subgraph can be found in the aforementioned works and 161 of the remaining 162 UniProt indexing nodes are not found in literature and therefore can be considered complete novel findings (Supplementary Table 6).

Then, we validate our findings against experimentally obtained results. Stukalov et al. [10] identified the human-virus interactomes of SARS-CoV and SARS-CoV-2 as consisting of 1801 interactions between SARS-CoV/SARS-CoV-2 viral proteins and 1086 human proteins (1082 of them can be mapped to the *human* Biomine database). Li et al. [11] experimentally identified 295 SARS-CoV-2 virus-host protein interactions (between SARS-CoV-2 viral proteins and 286 human proteins) and found potential molecular mechanism for SARS-CoV-2-induced cytokine storm. We are able to map 284 of the human proteins to the *human* Biomine database. To validate the robustness of our discovered dense sub-communities, we re-did our analysis and considered their discovered PPIs as additional ground truth. Our experiments indicate that by integrating more ground truth nodes from these two studies, our final extended network has a nearly identical structure as the presented ones in our paper and with exactly the same 3625 nodes. The main change is that more nodes in our final merged subgraph are being validated and the total number of generated hypotheses (nodes in the extended network that do not belong to the integrated experimentally validated results) has decreased, indicating that our identified dense sub-communities are robust enough. Besides validating the network, we also validated the biological stories of our findings. For example, SRC, ABL1, and JAK1/2 are generated hypotheses and we identified SRC, ABL1 and JAK2 to be important hub genes in previous section. In [10], JAK1 and JAK2 have been experimentally validated to interact with SARS-CoV and SARS-CoV-2 viral proteins. Similarly, we identified HSPA4, XPO1, CDK4, KIT to be among the high quality (most likely to be true) hypotheses in Table 3. Li et al. [11] have experimentally validated that HSPA4 interacts with SARS-CoV-2 N protein, XPO1 interacts with SARS-CoV-2 NSP8 protein, CDK4 interacts with SARS-CoV-2 NSP10 protein, and KIT interacts with SARS-CoV-2 ORF3a protein.

## Discussion

Using the rich Biomine database, we extended the SARS-CoV-2-human protein-protein interaction (PPI) network identified by Gordon et al. [8] and also integrated the discovered lists of pro/anti-SARS-CoV-2 genes by Wei et al. [9]. Through the proposed three-stage analysis pipeline, we were able to filter the large extended network, uncover dense sub-communities, and therefore generate research hypotheses related to COVID-19 by identifying dense cores in the Biomine database that have as many same nodes as the integrated experimentally validated results.

There are two fundamental assumptions in this work. Firstly, to extend the experimentally validated results (we mainly focus on the PPI network), we assume the results by Gordon et al. and Wei et al. to be correct. Secondly, to identify a small list of important proteins from a large network, we assume sub-communities of nodes (that are highly-connected with each other and well connected with experimentally validated nodes) to be more important than other nodes. In the literature, highly connected nodes such as hotspots or hub genes [10, 91, 92] have received substantial attention. Our analysis targets ‘high-activity sub-network’ and can be considered as an extension to hotspot (i.e. a cluster of connected hotspots, which is a higher level structure and is harder to detect). This is one of the novel contributions in this work.

In the first step of our proposed pipeline, we have two data-screening sub-steps added; this design is based on our assumption and will enable the peeling algorithm (PA) to focus more on cores with high activities. Furthermore, by focusing on dense cores, the PA can run faster due to a decrease in the input dataset size.

It is worth noting that when the network contains more than one connected component after the two-stage data screening, our proposed pipeline will produce the result even faster since the network complexity will be further reduced and we can run PA in parallel for each connected component. But in Biomine, the network is connected after screening and we were unable to run PA in parallel.

Based on the aforementioned assumption, the Biomine database, and the proposed analysis pipeline, we quickly identified sub-communities in the extended PPI network that have high activities and we discovered novel diseases, genes, and proteins that could potentially relate to COVID-19 as research hypotheses. The generated hypotheses (Supplementary Table 5) provide candidates for follow-up work to validate.

Kumar et al. [20] also employed a similar graph decomposition concept on a host-viral network. The difference between their analysis approach and ours is that they started with a deterministic graph and added edge weight later while we started with probabilistic graph decomposition which is more challenging. Another notable difference is they utilized a graph decomposition method called weighted *k-shell*. In contrast to our *k-core* decomposition, *k-shell* asks for all the nodes in the subgraph to have coreness precisely equal to *k* while *k-core* asks for all nodes’ coreness to be at least *k*. *k-core* has the advantage of connecting different subsets of nodes in the network and helps discover missing links between them. Additionally, when we perform pathway and process enrichment analysis on the subgraph, we also use the subset of nodes with the same coreness, which by definition is the *k-shell* of the subgraph.

There are three user-defined thresholds in the first and second steps of our proposed analysis pipeline: threshold based on degree expectation, threshold based on *η-degree* lower-bounds, and *η*. These thresholds can be changed based on users’ specific needs. The first two thresholds are used to filter out nodes with very low connectivity, which cannot be members of dense sub-communities in the network. By filtering out low connectivity nodes, we reduce the network volume and speed up the analysis. So, the first threshold should be a number much smaller than the coreness of nodes in the dense sub-network. In our analysis, we set the threshold as 5, and we are interested in the dense sub-network with coreness ≥ 69, hence threshold 5 is a conservative choice. As long as the first threshold is set relatively far away from the coreness in our final dense subgraph, their value only affect the computation speed, but not affect the results. The user could use a larger thresholds to increase speed, if they are ok to take more risk of missing important nodes. Similar rule also applies to the second threshold. For the third threshold, *η*, its choice is rather subjective, but we found the results of our analysis are robust to the choice of its value. Figure S3 in Supplementary Methods and Results shows the results of different settings of *η* (0.3, 0.5, 0.7) to be very consistent (nodes’ coreness under different *η* values are in almost perfect linear relationship).

For determining the merged subgraph, we need to pick an optimal threshold of coreness that will balance the denseness and the complexity of the graph. We observed that core 69 and 77 are both dense, frequently appeared, and contains more information. Compared to core 77, we preferred core 69 as we want to include more nodes into the subgraph. In another hand, it is not practical to involve too many nodes which results in generating a vast amount of hypotheses to validate. Therefore after comparing with coreness 77 and other core number in Table 1, we chose coreness 69 as the final optimal threshold.

We did a literature review for most of the genes detected in our subgraph and listed top genes in Table 3. Take entries with *nConnect* ≥ 110 as an example, 7 genes received literature support (we will discuss the tyrosine-related ones together in the next paragraph). The other 15 genes with *nConnect* ≥ 110 might be strong candidates with high priority and worth for future validation. For GAPDH (Glyceraldehyde-3-phosphate dehydrogenase), Zheng et al. [42] obtained potential COVID-19 effector targets of Chinese medicine Xuebijing (XBJ) which was used in treating mild cases of COVID-19 patients, and GAPDH is found to be one of the key targets. Taniguchi-Ponciano et al. [43] identified HIF1*α* as a potential marker for COVID-19 severity and GAPDH is found to be among the expressed HIF1*α* responsive genes. Ebrahimi et al. [44] found inhibiting GAPDH in patients with degenerated innate immunity can potentially help in treating COVID-19. For DICER1 (Endoribonuclease Dicer), Mu et al. [45] found that SARS-CoV-2 N protein prevent Endoribonuclease Dicer from the recognition and cleavage of virus-derived dsRNA. For GSK3B (Glycogen synthase kinase-3 beta), Liu et al [50] investigated COVID-19 traditional Chinese medicine treatment ShenFuHuang formula and identified GSK3B as the medicine’s potential drug target. Khalil [51] and Nowak et al. [52] hypothesized lithium chloride (directly inhibits GSK3B) to be a potential treatment for COVID-19 due to its inhibition effect on other members of the Coronaviridae (CoV) family. Embi et al. [53] found chloroquine treatment (preventing SARS-CoV-2 to fusion with the host cell membrane) results in the inhibition of GSK3B.

For tyrosine-related proteins in the merged subgraph and their associations with COVID-19, many tyrosine kinase inhibitors have been identified to inhibit the SARS-CoV-2 virus. Cagno et al. [93] found three ABL tyrosine-protein kinase inhibitors imatinib, dasatinib, and nilotinib to exert inhibitory activity against SARS-CoV-2, Alijotas-Reig et al. [94] found two JAK tyrosine-protein kinase ruxolitinib and baricitinib to be useful in treating the COVID-19 induced systemic hyperinflammatory response (cytokine storm). Wu et al. and Seif et al. [56, 95] found another JAK inhibitor fedratinib to mitigate the serious conditions in COVID-19 patients. We identified SRC, JAK1/2, ABL1/2 as hub nodes in Figure 5. For SRC (Proto-oncogene tyrosine-protein kinase SRC), in Lin et al. [46], ibrutinib is found to block SRC family kinases, which might reduce viral entry as well as the inflammatory cytokine response in the lungs. Morenikeji et al. [47] identified SRC to be one of the genes associated with Bovine coronavirus and by implication, other coronaviruses. Tiwari et al. [48] found SRC to play a vital role in SARS-CoV-2 infection related pathways. Xie et al. [49] found SRC participates in cytokines storm in patients with obesity which could lead to negative outcomes when infected with SARS-CoV-2. Additionally, many works in literature have targeted JAK1/2 in the hope to treat or prevent COVID-19. Shi et al. [79] found decreasing of lymphocyte in patients with COVID-19 correlated with low expression of JAK1-STAT5 signaling pathway. Zhang et al. [58] found that by suppressing JAK1/2 using baricitinib, several cytokines signals inciting inflammation will be inhibited. Others have also suggested using drugs that inhibit JAK1/2 to help patients with COVID-19 [59, 60, 61, 62, 63, 80, 81, 82]. As for ABL1/2, Abruzzese et al. [55] have suggested a possibility that patients treated with BCR-ABL tyrosine kinase inhibitors may be protected from the virus infection.

## Conclusion

With the SARS-CoV-2 outbreak being declared as a pandemic by the World Health Organization (WHO) [96], researchers around the globe are shifting their focus to COVID-19. In this work, we are especially interested in the protein-protein interaction (PPI) network between SARS-CoV-2 proteins and human proteins identified by Gordon et al. [8] and the lists of pro/anti-SARS-CoV-2 genes discovered by Wei et al. [9], and our focus mainly lies in extending the network to generate biological hypotheses for further validation. To achieve that, we connect the single experiment derived PPI network with the large Biomine database, also integrate the findings of Wei et al., and aim to locate sub-communities with high activities in the extended network. We propose a data analysis pipeline based on a graph peeling algorithm (PA) that enabled us to compute core decomposition efficiently. We select dense cores in the Biomine database that overlap most with the integrated experimentally validated results. The dense subgraph is the resulting extended network and nodes not belonging to the integrated experimentally validated results in the subgraph are generated hypotheses. We then evaluate the selected subgraph in three contexts: we performed literature validation for uncovered virus targeting genes and proteins and found genes that have already been validated by others on their relationships to COVID-19; we carried out gene ontology over-representation test on the subgraph and found underlying enriched terms related to viral replication, viral pathogenesis, cytokine storm, etc.; we also searched for literature support on the identified tissues and diseases related to COVID-19 and found the possibility of drug repurposing for COVID-19 treatment. To further assign priorities to the generated hypotheses, we sorted all UniProt indexing nodes in the subgraph by their connections to the integrated experimentally validated nodes. The top ranking nodes (Table 3) in the list have a high proportion of literature validated nodes (for instance, GAPDH, DICER1, GSK3B, UBC, HSP90AA1, HSPA8, and tyrosine-protein kinase SRC, JAK1, JAK2, ABL1, etc.), we deem the rest non-validated nodes in the table as high quality hypotheses.

## Code availability

Complete methods for data preprocessing and peeling algorithm (PA) from analysis pipeline can be found on GitHub (https://github.com/ubcxzhang/COVIDnetwork).

## Supplementary materials

The interactive version of Figure 6-(a), Supplementary Report 1, and Supplementary Tables 1-6 can be found on GitHub (https://github.com/ubcxzhang/COVIDnetwork/-tree/master/Supplementary_materials).

## Author contributions statement

X.S. and X.Z. contributed to the study concept and design. Y.G. and F. E. contributed to the acquisition of the datasets, data processing and data analysis. Y.G. wrote the first draft with input from X.S. All authors have developed drafts of the manuscript and approved the final draft of the manuscript.

## Acknowledgments

The authors thank the anonymous reviewers for their valuable suggestions. We acknowledge Compute Canada for providing the computational resources. X.S. is supported by the National Research Council Canada through the Artificial Intelligence for Design program.

**Yang Guo** is a master’s student in the Department of Mathematics and Statistics at the University of Victoria, Canada. His research interests include bioinformatics and data mining.

**Fatemeh Esfahani** is a Ph.D. candidate in Computer Science at University of Victoria, Canada. Her research interests include graph algorithms, social network analytics, and data mining.

**Xiaojian Shao** is a research officer at Digital Technologies Research Centre, National Research Council Canada. He has research interests in bioinformatics, computational epigenomics and machine learning.

**Venkatesh Srinivasan** is a professor in the Department of Computer Science at the University of Victoria, Canada. His current research interests are in algorithms for big data, data mining, and data privacy.

**Alex Thomo** is a professor in the Department of Computer Science at the University of Victoria, Canada. His research interests include databases, algorithms for big graphs, social networks, and data mining.

**Li Xing** is an assistant professor in the Department of Mathematics and Statistics at the University of Saskatchewan, Canada. Her expertise is in biostatistics and statistical machine learning.

**Xuekui Zhang** is a Canada Research Chair in Biostatistics and Bioinformatics and an assistant professor in the Department of Mathematics and Statistics at the University of Victoria, Canada. His expertise is in biostatistics, bioinformatics, and statistical machine learning.

## Footnotes

1 332 viral-interacting human proteins and an additional human protein that interact with the viral-interacting human proteins

